# Design and validation of a novel portable electrode holder for motor-related EEG measurement

**DOI:** 10.64898/2026.03.02.705772

**Authors:** Mori Fukuda, Masaaki Hayashi, Seitaro Iwama, Junichi Ushiba

**Affiliations:** School of Fundamental Science and Technology, Graduate School of Keio University, Kanagawa, Japan; Department of Biosciences and informatics, Faculty of Science and Technology, Keio University, Kanagawa, Japan

**Keywords:** electroencephalogram, brain-computer interface, electrode placement, mobile EEG, upper limb motor function

## Abstract

**Objective:** A simplified headset to measure electroencephalography (EEG) from sensorimotor areas is necessary to monitor motor-related neural responses accurately in real-world environments. The aim of the present study was to design and validate a novel, easy-to-use, and reliable EEG electrode holder that enables positioning of the EEG electrodes directly over the sensorimotor cortex, while enabling flexibility in relation to varying head size.

**Approach:** The spatial distribution of motor-related EEG activity was estimated using a dataset of high-density EEG (HD-EEG) from 82 participants. International databases of head shape were analyzed to determine dimensional requirements for a headphone-shaped holder. The proposed headset was validated by comparing its recordings with those obtained with a commercially available HD-EEG (Experiment 1) and by computing the actual position and recorded EEG motor-related activity obtained using the proposed holder (Experiment 2).

**Main results:** The estimated distance between the measurement electrode and the top of the head, used as a design requirement for the proposed headset, ranged from 56.2 to 80.0 mm. Experiment 1 showed that the center frequencies of alpha-band recorded by the proposed and by the HD-EEG headsets were highly correlated (r=0.97). Experiment 2 showed that, by using the proposed headset, the actual placement of electrodes was within 8 mm from the ideal positions. Moreover, the experiment showed consistent results in terms of task-, location-, and frequency-specific modulation of sensorimotor activities in the alpha-band. For example, significant contralateral motor-related event-related desynchronization in the alpha-band, and significant alpha-band power increase during the eyes closed condition, namely event-related synchronization.

**Significance:** The proposed electrode holder is easy to use and adjustable to compensate for varying head size and it may enable reliable measurement of motor-related EEG. It could support practical application of motor-related EEG acquisition in real-world contexts in several applications including sports, rehabilitation, and artistic performance.

## 1. Introduction

Scalp electroencephalogram (EEG) is one of the most commonly used neurophysiological measurements as it can be used for assessing brain function in a noninvasive way and with excellent temporal resolution. These characteristics have supported the development of EEG-based brain-computer interfaces (BCI) for various purposes. Examples of EEG-based BCI applications include, but are not limited to: neurofeedback training for post-stroke upper-limb rehabilitation [1], neuromarketing [2,3], neuronal cytology [4], neuroeducation [5], sleep monitoring [6], sensory evaluation [7], and driver alert detection [8]. EEG-based applications are also widely adopted in experimental research for example for neural monitoring and closed-loop neurofeedback [9].

In recent years, a variety of studies have provided a growing body of evidence about the effectiveness of BCI applications for restoring upper limb motor-related neural activity, for example for post-stroke rehabilitation [1,10–13]. BCI-based rehabilitation is based on up-regulation of the excitability of the sensorimotor cortex (SM1) through the interplay of human brain activity and feedback based on recorded neural activity from BCI. Recently, several meta-analyses have demonstrated the effectiveness of BCI for post-stroke upper-limb rehabilitation [1,10,11]. Moreover, BCI can also be used to control assistive prosthesis combined with electromyography and robotic devices [12,14]. In these applications, EEG signals are measured in SM1 which plays a dominant role in motor execution [15,16]. The effectiveness of BCI, specifically in terms of neurofeedback training targeting motor-related areas, has been demonstrated even in healthy individuals in a meta-analysis of 33 studies, including 13 studies that employed EEG-based BCI focusing on SM1 activities [13].

To increase accessibility of BCI applications for motor rehabilitation, there is a need for portable EEG setups able to effectively and accurately monitor SM1 activity therefore supporting interventions across various real-world contexts. So far, several user-friendly headsets featuring different electrode configurations have been developed to streamline the BCI setup process. Examples include the “MuseTM” system by InteraXon (Toronto, Ontario, Canada) designed to support meditation, and the “NextMind” by Snap (Santa Monica, CA, USA). These EEG headsets were deployed for monitoring and/or modulation of brain activity in specific regions of interest (ROIs). For example, the occipital visual cortex for BCIs based on visual evoked potentials [17–19], or the frontal region for investigating attention, emotions, and cognition [20–22]. However, to date there are no commercially available setups targeting SM1 in a sufficiently specific manner.

Precise placement of electrodes is of paramount importance for reliable interpretation of signals from SM1 because of somatotopy in this area, i.e., localized functional dominance corresponding to specific body parts. Due to this characteristic, electrodes over SM1 provide different responses depending on the activated body parts. Moreover, EEG recordings from SM1 are challenging also due to interindividual variability in head size. In conventional EEG measurements, adaptation to head size is generally addressed using electrode placement frameworks that utilize standardized ratios derived from cranial landmarks (e.g., the 10-20 method) [23]. However, the simpler setup of portable EEG systems, which is based on fixed electrode distances, does not allow adaptation to varying head size. To support wider use of portable EEG systems targeting SM1 for real-world application of BCI in motor rehabilitation, it is crucial to design and validate reliable, portable headsets for motor-related EEG measurements, able to provide accurate signal measurements for a significant portion of the population, overcoming the abovementioned issues related to spatial spread of neural activity and interindividual variability in head size [9,24–26].

The aim of this study was to design and validate a novel electrode holder to detect motor-related activity. The study was conceived as a three-step process:

1. evaluation of the quality of motor task event-related desynchronization (ERD) of the sensorimotor rhythm (SMR) assessed using an available EEG database. The ERD used in this study is a modulation found in SMR, which is attenuated by motor output due to neural desynchronization in the sensorimotor cortex [27–29]. Since ERD is thought to be observed mainly due to the interplay between the sensory and motor cortices [28] and is sensitive to motor execution and motor imagery, conventional BCIs can successfully detect the presence/absence of motor output [30–33]. In this study, the spatial distribution of ERD was analyzed from an open 129-channels high-density EEG (HD-EEG) dataset of 82 participants performing right-hand motor task [34] by analyzing fluctuations of the alpha-band, which is known to reflect cortical function.
2. Definition of requirements for electrode placement that allow ERD to be observed using the anthropometric database [35–37]. Specifically, using available data about the spatial extent and the head size from three countries (United States [35], Japan [36], and China [37]), we calculated the conditions required for the holder dimensions (i.e., the distances between Cz-C3 and Cz-C4) that could be used for ERD measurements in the largest population possible by considering the maximum and minimum dimensions of the head.
3. Validation of ERD during a motor execution task assessed using a portable electrode holder reflecting the requirements defined in (2). Specifically, two experiments were conducted: the first experiment was performed with the newly designed electrode holder only, and the second experiment was performed using both the proposed electrode holder and an HD-EEG net-cap.

## 2. Methods

### 2.1. Spatial characteristics of EEG-SMR-ERD

The spatial distribution of SMR ERD in HD-EEG during a unilateral hand movement task was computed using a publicly available dataset [19]. After preprocessing and noise removal, spatial regions showing similar EEG responses were determined based on two criteria. First, regions where the magnitude of the response was close to the one measured at the center of the ROI were identified. Second, regions where the temporal waveform of the response was highly correlated to the one measured at the center of the ROI were identified.

#### 2.1.1. Datasets & preprocessing

The EEG database here used includes EEG data from three different studies, for a total of 82 participants (74 males and 8 females, mean age 21.08 ±1.4, all right-handed). Participants repeatedly performed a resting task and a motor imagery (MI) task of right-hand finger extension (i.e. contraction in extensor digitorum communis), while EEG was recorded using a 128-channel EEG cap (Geodesic Sensor Net, Electrical Geodesics Incorporated, Eugene, OR, USA) at a sampling rate of 1000 Hz. **Figure 1**(a) shows the experimental protocol of each experiment, consisting of three paradigms (A, B, and C) involving 30, 30, and 22 participants, respectively. The three paradigms included a fixed number of repetitions of a basic trial which included a sequence of rest, an interval of preparation, the MI task, and an additional interval before the next trial. The three paradigms had varying number of repetitions and varying duration of the rest, preparation, and MI intervals. Since these experimental protocols were designed to train MI with BCI feedback, only the first block of trials was used for analysis to avoid the influence of training effects.

**Figure 1.**
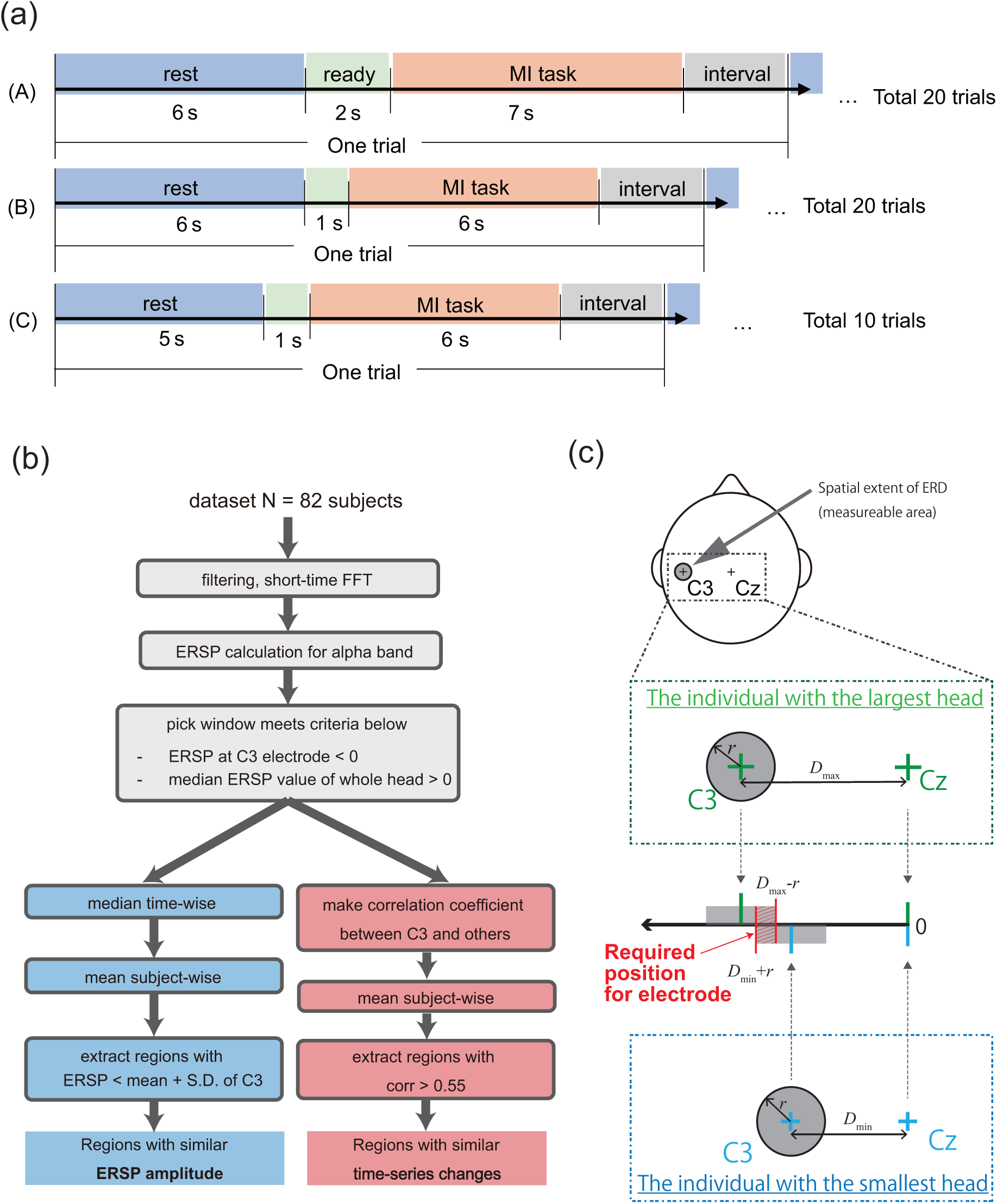
Derivation of the required electrode placement based on the dataset. (a) Schematic representation of the experimental protocol of the study dataset. Datasets from paradigms A, B, and C include recordings from 30, 30, and 22 participants, respectively. (b) Summary of the signal processing pipeline. The blue and red boxes report the steps related to the identification of regions with similar ERD amplitude and temporal waveform, respectively, compared to the C3 electrode. (c) Identification of the range for Cz electrode position based on spatial region of ERD detection. ***D*_min_** and ***D*_max_** were calculated from anthropometrical databases based on minimum and maximum head dimensions, respectively. The radius ***r*** was estimated from the EEG database through the processing pipeline shown in panel (b).

A schematic representation of the analysis pipeline is shown in Figure 1(b). The acquired EEG signals were de-noised with a sixth order Butterworth bandpass filter with a passband of 3-40 Hz and a 50 Hz notch filter (implemented as a sixth order Butterworth band-stop filter with stop band of 49-51 Hz). Then, the spectrum was obtained every 100 milliseconds using the Short-time Fourier Transform (STFT) with a 1 second Hanning sliding window, resulting in 90% overlap. The event-related spectral perturbation (ERSP) for a given electrode *e* was obtained using the following formulas:

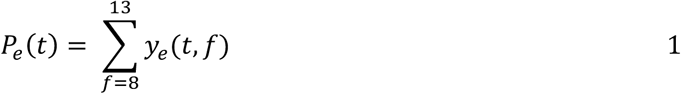

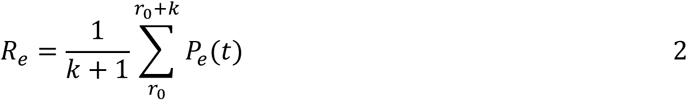

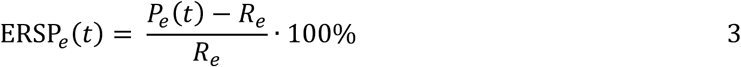

where *y_e_*(*t, f*) is the power spectral density computed at frequency *f* and time *t*, *P_e_*(*t*) is the alpha-band amplitude, *R_e_* is the mean alpha-band amplitude in the resting-state epoch [*r*_0_, *r*_0_ + *k*]. Negative values of ERSP indicate occurrence of ERD and enable estimation of its depth [27,38].

To obtain the spatial distribution of motion-related ERD in sensory-motor area, ERSP segments that satisfied the following criteria were extracted: (1) ERSP_C3_(*t*) ≤ 0 (where ERSP_C3_ is ERSP at C3 electrode), i.e. ERD detected at C3 electrode; (2) median ERSP > 0 for all electrodes, i.e., ERD did not occur.

#### 2.1.2. Analysis of spatiotemporal characteristics of SMR-ERD

To identify a set of electrodes that show similar ERSP trends compared to the C3 electrode, we evaluated two key aspects: ERSP amplitude and temporal synchrony. The rationale is to determine a spatial distribution of electrodes on the scalp that delineates the region in which the ERD can be effectively observed.

Electrodes with mean ERSP amplitude similar to that recorded at the C3 electrode, which is defined as the center of the ROI, were identified following the procedure summarized in the blue boxes in **Figure 1(b)**. Specifically, for each trial, we computed the median ERSP over time to limit the effects of noise and we subsequently averaged the median ERSP across all participants and all trials in the block. The region displaying ERSP negative amplitudes higher, in absolute value, than a specified threshold was defined as the spatial region for ERD detection. The threshold was determined as the mean of the ERSP values at C3 plus one standard deviation, in order to encompass about 84% of ERSP values, assuming a normal distribution of ERSPs at the center of the ROI. The normality of the ERSP distribution at C3 was confirmed by the one-sample Kolmogorov-Smirnov test. (statistical significance was set at p < 0.05).

Electrodes with temporal ERSP patterns similar to those recorded at C3 were identified following the procedure indicated in the red boxes in **Figure 1(b)**. Specifically, time-series Pearson’s correlation coefficients for ERSP between each electrode and C3 were computed. The spatial region for ERD detection was defined as the area in which the correlation coefficient was higher than 0.55, indicating a moderate positive correlation [39,40].

### 2.2. Estimation of optimal electrode positions

To account for interindividual variations in human head dimensions, the design of the headset was defined based on the ERD measurable range, estimated as described in section 2.1. Given the focus on sensorimotor cortex, a headphone-type enclosure was chosen. Accordingly, we assessed dimensional values related to the Bitragion Coronal Arc length.

To gather sufficiently generalizable head dimension data, we analyzed human body dimension databases from three countries: the United States [35], Japan [36], and China [37]. Using the spatial range determined following the procedure described in section 2.1, we defined the distance between C3 and Cz electrodes, denoted as “*D*”, to fall within the following specified range:

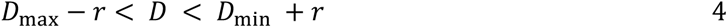

where the *r* is the spatial length of ERD (i.e., the gray region shown in Figure 1(c)) on the arc, *D*_max_ is the distance between Cz and C3 electrode in the largest head, and *D*_min_ is the distance between Cz and C3 electrode in the smallest head. As a result, the electrode holder was designed in a way that the slider mechanism comes around the ear, allowing the headset to be operated with straightforward positioning procedures.

The headset was equipped with an application-specific integrated circuit-based analog circuit and microprocessor derived from an existing electroencephalograph: the AE-120 (Nihon Kohden Corp., Tokyo, Japan). EEGs were derived in a monopolar manner and processed with ×1200 amplification and 0.21 to 199 Hz analog band-pass filtering on circuit. The processed EEG signals were digitized at 200 Hz with 12 bits resolution. In the measurement described below, EEGs were then transmitted to a laptop using a Bluetooth 3.0 wireless protocol. A 3 to 35 Hz bandpass filter with a 50 Hz notch was used to reduce noise contamination. A MacBook Air (M1 2020) was used for storing the streamed signals and for delivering the protocol to participants.

### 2.3. Validation of the proposed headset during a motor execution task

#### 2.3.1. Experiment 1: Simultaneous measurement with a HD-EEG headset

##### 2.3.1.1. Experiment design

In this experiment, we measured EEG modulation during a motor task while wearing both the proposed electrode holder and an existing HD-EEG. Participants performed the task repeatedly for 80 trials (10 trials/block * 8 blocks) while EEG signals were recorded. In each trial, the instructions were presented in the following order: ready1, rest, ready2, task, interval. Participants pressed a key at any point during the interval epoch to start the ready1 epoch, which was followed by a two-seconds countdown and six-seconds rest (as shown in **Figure 2(a)**). Then, in the ready2 epoch, the subject was presented with two-seconds countdown and six-seconds task. The task consisted in motor execution (ME) of a right finger extension. These cues were repeated ten times, resulting in one block. At the beginning of each block, the impedance was checked to be below 50 kΩ before starting the measurement.

**Figure 2.**
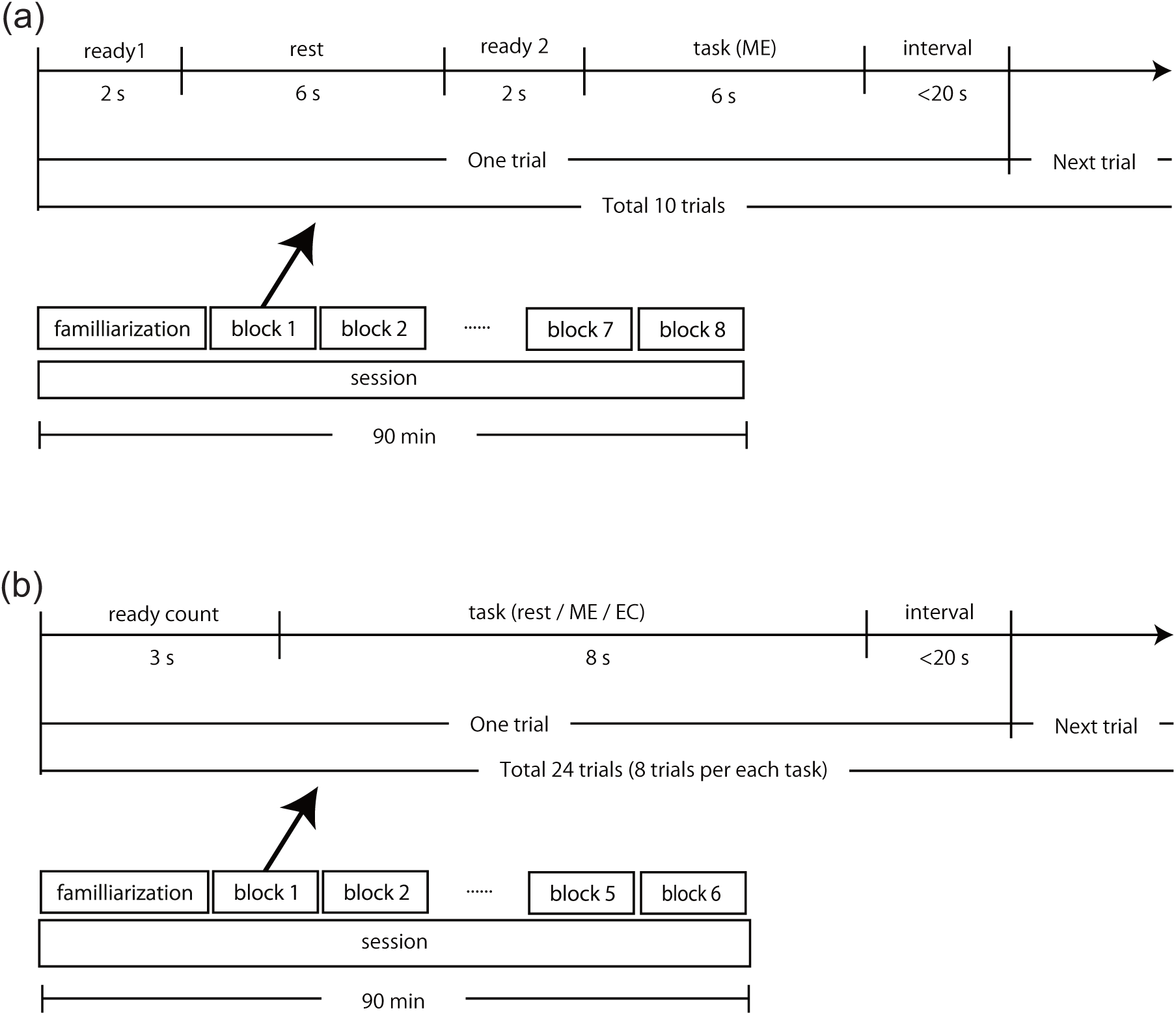
Experimental protocols. (a) Experimental protocol of the simultaneous EEG recordings using the proposed electrode holder and the HD-EEG net cap. The task consisted in motor execution of the right hand. (b) Experimental protocol of EEG recordings using a proposed electrode holder alone. The task was randomly assigned to rest, rest with eyes closed (EC), or motor execution (ME).

The HD-EEG data were recorded using 128 channel counts from the Hydrogel Geodesic Sensor Net 130 system (Electrical Geodesics Incorporated [EGI], Eugene, OR, USA) with a sampling rate of 1 kHz in a quiet room. The proposed headset was attached over the net cap by the experimenter before each session.

##### 2.3.1.2. Participants

Ten healthy individuals (females: 3, age: 20-27 years old) participated in the experiment. Participants with mental retardation, head injury, loss of consciousness, or other neurological disorders that might have affected the experiment were excluded. The study was conducted in accordance with the Declaration of Helsinki. The experimental procedure was approved by the Ethics Committee of the Faculty of Science and Technology, Keio University (Number: 2022-120). Written informed consent was obtained from all participants before the experiments.

##### 2.3.1.3. Data analysis

###### Preprocessing

The EEG signal was filtered between 3 to 35 Hz with a fourth-order Butterworth bandpass filter and a 50 Hz notch filter. Subsequently, the EEG signals of all electrodes were subtracted by the signal from Cz to enable the detection of local alterations within the sensory-motor cortex.

###### SMR-ERD during motor execution task

The amplitude of individual alpha frequency (IAF) was extracted by using the STFT as described in section 2.1.1 and the ERD in the motor cortex during the ME task was addressed by analyzing the amplitude of IAF. The amplitude of the IAF was computed in a 3Hz band centered around the highest peak in the power spectrum extracted in the range of 8-13 Hz from the EEG in the rest period. The ERD during ME was obtained by calculating the relative power to the average power computed during the resting epoch, as shown in Equations 1-3. From the ERD time series, the median value was computed and used as the estimate of ERD depth in the trial to limit the effect of amplitude fluctuations due to noise.

###### Frequency component analysis

The frequency spectra obtained with the STFT were fitted using the Fooof Algorithm [41]. This algorithm models the power spectrum as a combination of aperiodic signals and putative periodic oscillations. Spectral consistency between signals recorded using the proposed and the existing headset was evaluated using the Pearson’s correlation coefficient at the center frequencies of periodic oscillation components extracted by the Fooof Algorithm. A test of no correlation was performed to address significance of the estimated Pearson’s correlation coefficients (statistical significance was set at p < 0.05).

#### 2.3.2. Experiment 2: Stand-alone performance

##### 2.3.2.1. Experiment design

We used the proposed electrode holder to observe EEG modulation during motor tasks. Participants performed the task repeatedly for 144 trials (24 trials/block * 6 blocks) while EEG signals were recorded. The instructions were presented in the following order: ready, task, and interval. Participants initiated the ready epoch by pressing a key at any point during the interval epoch. During the ready epoch, a 3-second countdown was presented, followed by a 6-second task epoch, and finally the interval epoch. This sequence was repeated for 24 trials and encompassed four different tasks: rest, rest with eyes closed (EC), ME. These tasks were randomly cued 8 times each within the session. At the beginning of each session, the impedance was checked to be below 50 kΩ before starting the measurements. Some participants (ID 17-28) performed the tasks for 192 trials (32 trials/session * 6 sessions) including the MI task of right finger extension. Since the primary focus is on the observation of ERD induced by ME in a region-specific manner, in this study we did not evaluate the ERSP during MI task. While It is widely assumed that MI activates similar pathways as ME [42,43], and recognizing that no feedback on alpha amplitude was provided during any of the tasks, we assume no positive impact on the depth of ERD. Consequently, only the trials involving subjects’ ME are included in the analysis.

##### 2.3.2.2. Participants

Twenty-eight healthy volunteers (females: 14, age: 18-34 years) participated in this experiment. Participants with mental retardation, head injury, loss of consciousness, or other neurological disorders that might have affected the experiment were excluded. Those experienced in EEG measurements were also excluded. The study was conducted in accordance with the Declaration of Helsinki. The experimental procedure was approved by the Ethics Committee of the Faculty of Science and Technology, Keio University (Number: 2022-120). Written informed consent was obtained from all participants before the experiments.

##### 2.3.2.3. Data analysis

The acquired EEG signals were processed following the same procedure as in the first experiment, as described in Section 2.3.1. Similarly, ERDs were obtained following the procedure described in the first experiment (Section 2.3.1).

###### Electrode placement error

The electrode placement error in the proposed electrode holder was assessed by computing the distance between the ideal C3 (and C4 which is positioned symmetric with C3) position and the actual electrode position. This distance was manually determined through analysis of photographs taken immediately after the session, with a precision of 1 millimeter.

## 3. Results

### 3.1. Spatial distribution of ERD estimated from EEG dataset

The spatial distribution of the mean ERD estimated from the EEG dataset is shown in **Figure 3**(a), whereas the measured correlation with the signal from C3 electrode is shown in **Figure 3**(b). The cut-off threshold value for ERD amplitude, defined as the mean plus one standard deviation, was equal to 39.2%, which equals to 83.4th percentile of ERSP at C3. **Figure 3**(c) shows the spatial region where activity equivalent to that of electrode C3 was observed, based on criteria described in section 2.1. The figure shows that similar neural activities were observed within a circular region, with a radius of approximately 25 mm centered around C3. Moreover, a significant departure from a normal distribution was not confirmed through statistical testing (one-sample Kolmogorov-Smirnov test, k = 0.1047, *p* > 0.05) for ERSP at C3.

**Figure 3.**
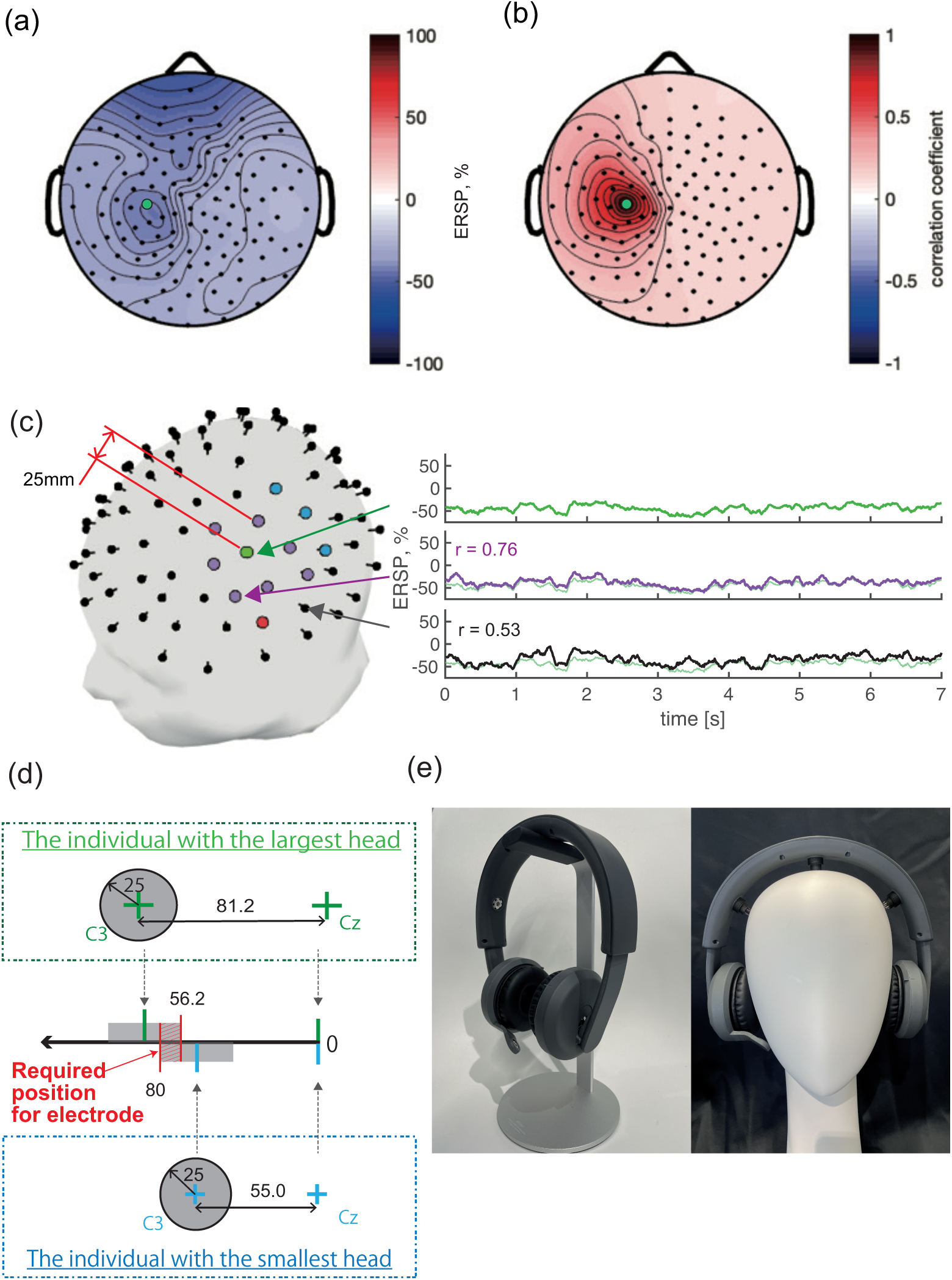
Spatial distribution of ERD, and electrode holder design. (a) Distribution of the mean depth of ERD during motor tasks estimated from the EEG dataset (N = 82 participants). The green dot shows the position of the C3 electrode. (b) Distribution of the correlation coefficients between ERSP measured from each electrode and from C3. (c) Distribution of electrodes showing similar activity and exemplary waveforms. Purple dots identify electrodes satisfying the two selection criteria: modulation deeper than mean + one standard deviation in C3, and correlation with C3 higher than 0.55. Blue dots show electrodes satisfying only the first criterion, whereas red dots show electrodes satisfying only the second criterion. (d) Estimated range of distance between C3 and Cz. The green marks show the required range for the subject with the largest head, and the blue marks show the required range for the subject with the smallest head. (e) Proposed headset developed based on the dimensional requirements shown in (d).

### 3.2. Dimensional requirements for the electrode holder

**Table 1** summarizes the head dimensions for each country and the resulting ideal Cz-C3 electrode distance. The maximum and minimum length of Bitragion Coronal Arc were equal to 406.0 mm and 275.0 mm, respectively. The estimated value of *r*, as defined in Equation (4), was equal to 25.0 mm based on results reported in Section 3.1. Accordingly, the estimated range of acceptable values for the Cz-C3 distance of the electrode holder was 56.2-80.0 mm (**Figure 3**(d)). The resulting electrode holder is shown in **Figure 3**(e).

**Table 1.** Estimated range of distance between C3 and Cz based on the maximum and minimum values of Bitragion Coronal Arc. Data in China[37], Japan[36], and the U.S[35].

| Country | Acceptable distance between C3 and Cz electrodes [mm] |  | Distance between C3 and Cz [mm] |  | Length of Bitragion Coronal Arc [mm] |  | Sample size<br>Male / Female | Age |
| --- | --- | --- | --- | --- | --- | --- | --- | --- |
|  | Min.* | Max.** | Smallest subject | Largest subject | Smallest subject | Largest subject |  |  |
| China | **56.2** | **80** | 55.0 | 81.2 | 275 | 406 | 2026 / 974 | 18 - 66 |
| Japan | 56 | 88.2 | 63.2 | 81.0 | 316 | 405 | 250 / 250 | 18 - 88 |
| US | 53.4 | 103.4 | 59.6 | 78.4 | 298 | 392 | 1774 / 2208 | 17 - 51 |

\* : C3-Cz distance of largest subject - measurable extent;
\*\* : C3-Cz distance of smallest subject + measurable extent

### 3.3. Validation of the proposed electrode holder

#### 3.3.1. Experiment 1: Simultaneous measurement with a HD- EEG headset

By using the proposed electrode holder, all the measurement electrodes were placed close to the target position of the existing net cap, regardless its size (**Table S1**). Approximate locations of electrodes on the proposed holder are shown in **Figure 4**(a), superimposed on the spatial pattern of the average ERSP obtained from the HD-EEG, averaged across the participants (N = 10). Results show that the proposed electrode holder placed the electrodes around C3 within a distance of about 20 mm, i.e., within the estimated radius of ERD.

**Figure 4.**
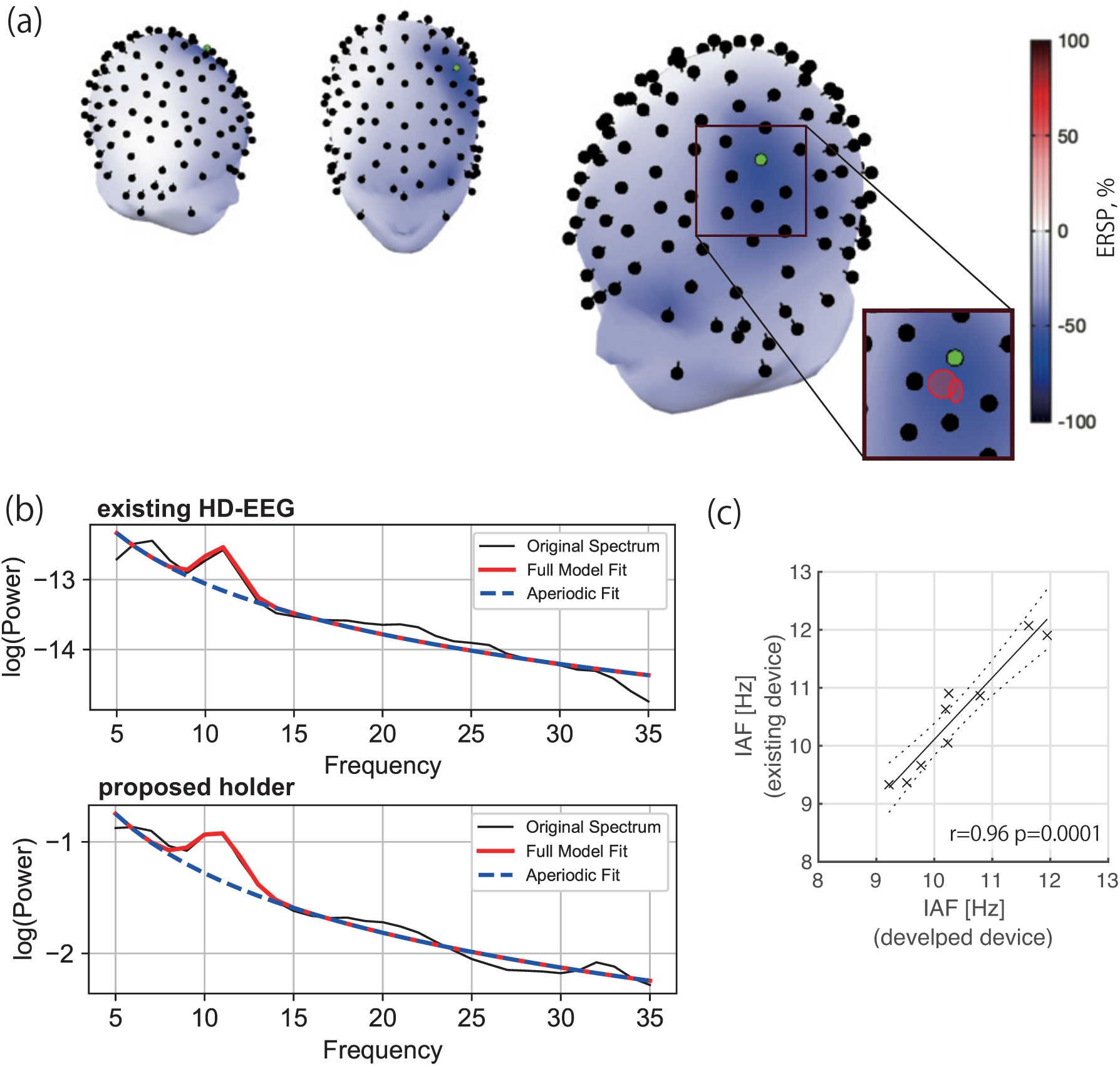
Comparison between the proposed headset and conventional HD-EEG. (a) Averaged ERSP spatial distribution and electrode placement. The green dot is the center of the measurement area (C3). The red shades are the regions where the electrodes were placed with the proposed electrode holder. Each region includes the actual position of electrodes in five participants. (b) PSD (black) and model fitted by Fooof algorithm (red: full model fit; blue: aperiodic fit) in one of the participants (ID=8). The PSD in the upper subplot is measured using the existing HD-EEG net cap, whereas the one on the lower subplot is measured using the proposed holder. (c) Scatter plot of the IAF extracted with the two headsets in the tested participants (*r*=0.97, *p*<0.05, CI=[0.83, 0.99], Pearson’s correlation coefficient, test of no correlation). N=9.

The power-spectral densities (PSD) acquired from both measurement systems at rest are depicted in **Figure 4**(b) (representative participant). A periodic component was obtained that peaked at the same frequency (11 Hz) by fitting in both devices. The R-square value of fitting was above 0.85 (mean±S.D. = 0.959±0.053) and the error is below 0.1 (0.061±0.029) in all conditions (participants and holder). After subtracting the aperiodic component, a peak was found in 9 out of 10 participants within the alpha frequency band (8-13 Hz). One of them did not show an alpha peak also in the case of the HD-EEG, and this absence may be due to an innate reason [44]. The central frequency of the component was defined as IAF. It was confirmed that IAFs detected by the existing net cap and the proposed holder are significantly correlated as shown in **Figure 4**(c) by Pearson’s correlation coefficient and test of no correlation (*r*=0.97, *p*<0.05, CI=[0.83, 0.99]). In one participant (ID=5), the IAF was not detected in both setups.

#### 3.3.2. Experiment 2: Stand-alone performance

The results of experiment 2 are shown in **Figure 5**. The placement errors between the ideal the actual positions of C3 and C4 were equal to -2 ± 6 mm. The distribution of placement errors is shown in **Figure 5**(a). In all participants, the placement error was lower than half of the acceptable range (i.e., 25 mm) defined in Section 3.1.

**Figure 5.**
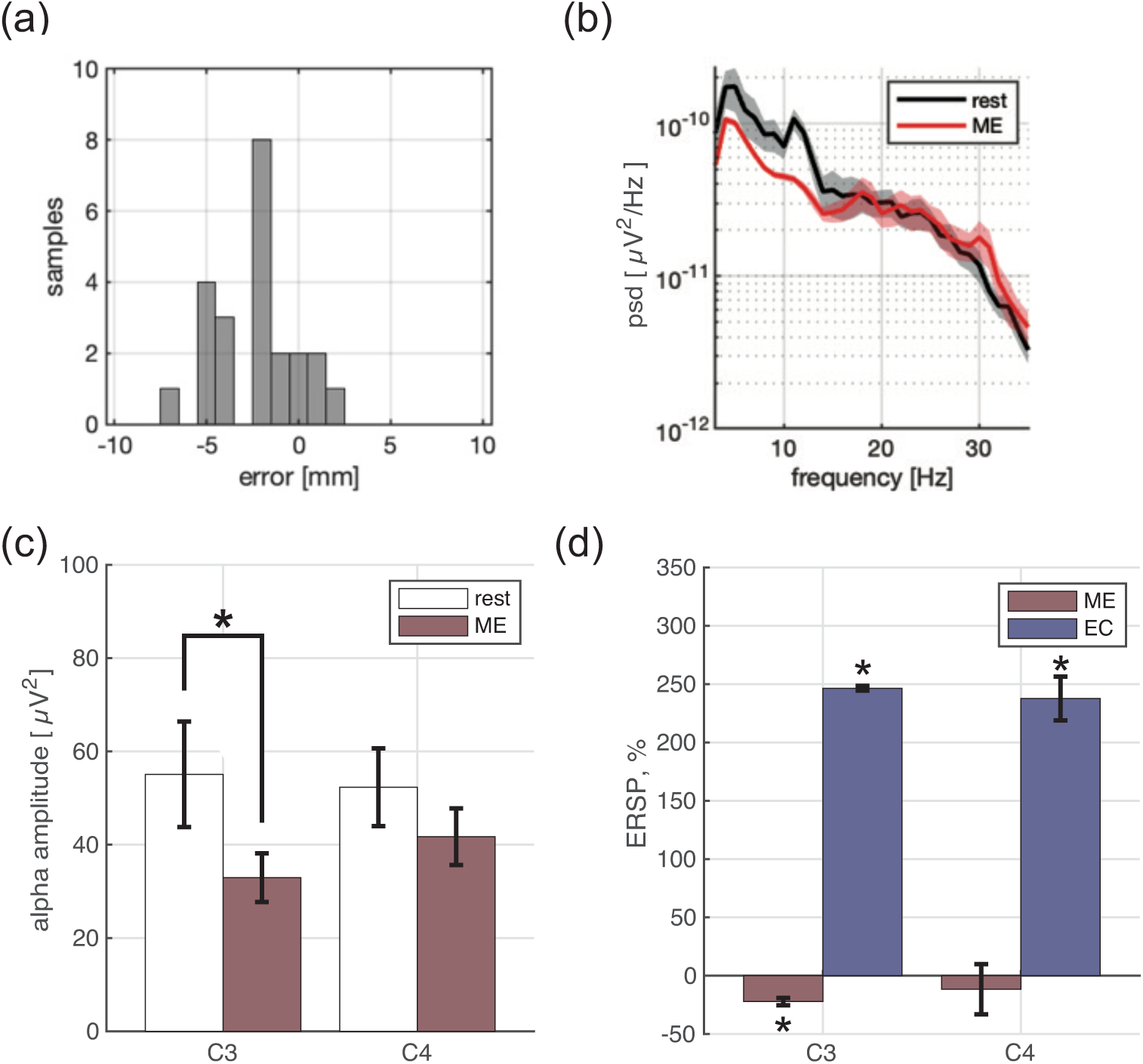
Sensor placement error and neural activity measured with the proposed holder alone. (a) Distribution of placement error for C3 and C4 electrodes using the proposed electrode holder. (b) Power spectral density (PSD, mean ± standard error) in C3 during rest (black line) and ME (red line). (c) Alpha-band amplitude modulation in C3 and C4 during ME compared to rest (two-sided paired t-test, *t*(12)=2.05, * *p*<0.05, Cohen’s *d*= 0.63, 95% CI = [-1.39, 45.68]). (d) Percent change in ERSP in ME epoch and EC epoch, compared to rest. Significant modulation was confirmed in C3 during ME and EC epoch, and in C4 during EC epoch (two-sided t-test, * *p*<0.05).

The power spectral densities (PSD) during rest and ME epochs in the C3 electrode are shown in **Figure 5**(b). Frequency-specific spectral modulation was confirmed around 11 Hz, which is known one of the central frequencies of the alpha-band [45]. **Figure 5**(c) shows the reduction in alpha-band amplitude for C3 and C4. Statistical analysis indicated that the observed reduction was significant only in C3 (paired t-test, *p*<0.05, Cohen’s *d*= 0.63, 95% CI = [-1.39, 45.68]). This finding is in line with the fact that ERD is usually observed just above the sensorimotor area contralateral to the limb executing the task [46]. The distributions of changes in ERDs in the ME and EC epochs compared to the rest epoch are depicted in **Figure 5**(d). These results indicate that the alpha-band amplitude increases when the eyes are closed. In addition, these results confirm that the spectral modulation was task-specific.

## 4. Discussions

In the present study, we designed and developed a novel electrode holder for movement-related EEG measurement. The design requirements were set based on movement-related EEG characteristics estimated from existing datasets and based on dimensional requirements derived from the international head dimension database. Motor event-related ERD was confirmed within a circular region with radius 25 mm centered around C3 (**Figure 3**). By putting this dimensional estimate in the context of real-world variability in head dimensions, a range of positions for electrode placement was determined in terms of distance between C3-Cz and C4-Cz, specifically a range of distance from 56.2 mm to 80.0 mm was obtained (**Table 1**, **Figure 3**(a-d)). The proposed holder has a headphone-like shape (**Figure 3**(e)) allowing intuitive alignment of the Cz position and it was designed to meet the electrode distance requirements defined above. The actual distance between Cz and C3 was approximately 75 mm and it varied depending on the head surface and height of the applied elastic electrode. In the proposed holder, the arm around the earpiece, away from the measurement electrode, can be slid to adjust the positioning to the individual head dimensions.

It is worth noting that the average value of Bitragion Coronal Arc computed from the international head dimension database may be used as an alternative method for electrode positioning. However, establishing the C3 electrode position (determined as 30% of the arc length) based on the average value computed using data from the three countries would result in a placement error up to 22.3 mm. This error was computed based on the minimum arc length (i.e., 275 mm) and the size of the sample utilized to calculate the global average of arc length (i.e., 349.42 mm) as indicated in Table 1. Specifically, the average arc lengths for each country are 354.09 mm (China), 343.93 mm (Japan), and 368.1 mm (U.S.), respectively. This error is within the acceptable range of 25 mm but the proposed electrode holder led to a considerably smaller error (up to 11.9 mm). Moreover, the proposed headset allows electrode navigation to the appropriate positions for all subjects without any individual adjustments, thus paving the way to EEG-based BCI technology for widespread use in real-world settings.

Specifically, the proposed electrode holder can place electrodes near the target positions (C3, C4, and Cz in the 10-20 method) in all participants, ensuring that all placements fell within the desired range (**Figure 3**, **Figure 5**). The position error was lower than 8 mm, that is a small error compared to other commercially available EEG headsets [9]. Validation of the headset versus simultaneous measurements from a HD-EEG net cap (as illustrated in **Figure 4**(c)) further supports the validity of the approach. In Experiment 1, the EEG measured with the proposed headset showed a 1/f-like background noise, in line with common observations [47] (**Figure 4**(b)), and the alpha frequency band amplitude decreased during the right-hand movement task. These findings demonstrate the task- and region-specific nature of these modulations (**Figure 5**(d)), supporting the potential use of the proposed electrode for high-quality, reliable EEG measurements.

In recent years, studies have shown that the amplitude and modulation of alpha frequency band may reflect cognitive function [48–51] and mental disorders [52,53]. Self-regulation training based on alpha-band feedback has been reported to improve motor function after stroke, supporting its potential therapeutic value [1,10]. In addition, the characteristics of the 1/f-like component have been increasingly recognized as a possible disease monitoring marker, specifically a macro-indicator of Excitation/Inhibition (E/I) balance [47]. These physiologically meaningful indices can be easily measured in humans and thus their use could support a range of clinical applications.

This study, by introducing a user-friendly and reliable EEG holder for hand motor-related EEG activity within the SM1 area, can potentially support motor function neuromonitoring while concurrently enhancing effectiveness of interventions, especially with regard to optimizing dosage [13]. Despite the availability of various portable headsets like MuseTM and NextMind, to the best of our knowledge, there is no product available in the market that specifically targets motor function in SM1 and has demonstrated consistent and reliable measurement capabilities in this regard. While some versatile general-purpose products like Quick-20m (Cognionics Inc. San Diego, USA) and Emotiv EPOC+ (Emotiv Systems Inc. San Francisco, USA) can encompass the SM1 region, there is a tradeoff between the number of electrodes and the ease of use of EEG devices [54]. Therefore, the development of purpose-specific electrode holders is imperative to meet the unique requirements of the target use cases.

It is worth noting that this study also provides a method to define the supported range of head dimensions for a headset, which can be useful for the design of future portable BCI. For instance, the method here used to define the desired electrode positioning range could be applied to define specific portable EEG headsets for children, for example for application in research related to cerebral palsy [55,56] or education [57,58]. Similarly, further adaptation may be needed to design headsets viable for use in individuals whose head dimensions fall outside the currently used anthropometric database.

### Limitations

This present study has limitations. For example, the study uses the inter-subject average of ERSP centered on the C3 electrode to define the target spatial region. Ideally, individualized spatial extents should be determined for each subject based on their specific neural characteristics. This is related to a number of considerations: (1) There is substantial variability in the shape of the cortical region with hand dominance among individuals [59–61]. (2) The center of the ROI, referred to as the “hot spot”, may not necessary coincide with the C3 position in the 10-20 method [62]. (3) There may exist correlations between the size of the dominant hand area and the dimensions of an individual’s head. Considering these factors, it would be important to develop a comprehensive HD-EEG dataset encompassing scalp dimensions, structural images, and hot spots, for example identified through Transcranial Magnetic Stimulation (TMS). These datasets could inform the design of devices that can accurately capture the distribution of EEG features in the real space, accounting for the regions common to the majority of individuals.

Another limitation is related to the fact that this study did not comprehensively address quality control after the setup, which is important for EEG measurement. Noise has several components, including noise from environmental or biological origin [45], which is often derived from the myopotentials of the eyes and facial muscles. Dealing with biological noise can be challenging as it requires guidance by the experimenter and attention from the person being measured. Although several noise reduction methods have been developed for post-processing [63–65], they can be difficult to apply, especially in portable headsets with a small number of electrodes, and measures need to be taken during measurement [54]. Future studies will need to develop an automated quality control system to ensure quality of recordings, particularly for simplified headsets such as the one here proposed.

## 5. Conclusion

Towards the final aim of realizing a portable EEG configuration capable of consistently and accurately measuring motor-related modulation within the SM1, this study introduced and validated a novel, portable EEG headset.

This innovative design enabled positioning of the electrodes precisely within the observation range of ERDs, demonstrated a signal structure similar to the one measured with the HD-EEG net cap, and confirmed the presence of task-specific, location-specific, and frequency-specific ERDs in relation to a motor task protocol.

The introduction of the proposed electrode holder holds significant promise for supporting the practical application of motor-related EEG acquisition in real-world contexts in a range of potential applications, for example sports, rehabilitation, and artistic performance.

## Acknowledgements

This work was supported by JST Moonshot R&D (grant number JPMJMS2012 to J.U). Junichi Ushiba is the Founder and Representative Director of the University Startup Company LIFESCAPES Inc. involved in the research, development, and sales of rehabilitation devices, including brain-computer interfaces. He receives a salary from LIFESCAPES Inc. and holds shares in LIFESCAPES Inc. S. Iwama is currently engaged in a contractual arrangement with the company for the provision of services. M. Fukuda and M. Hayashi is employed by the company.

---

**Table S1.**
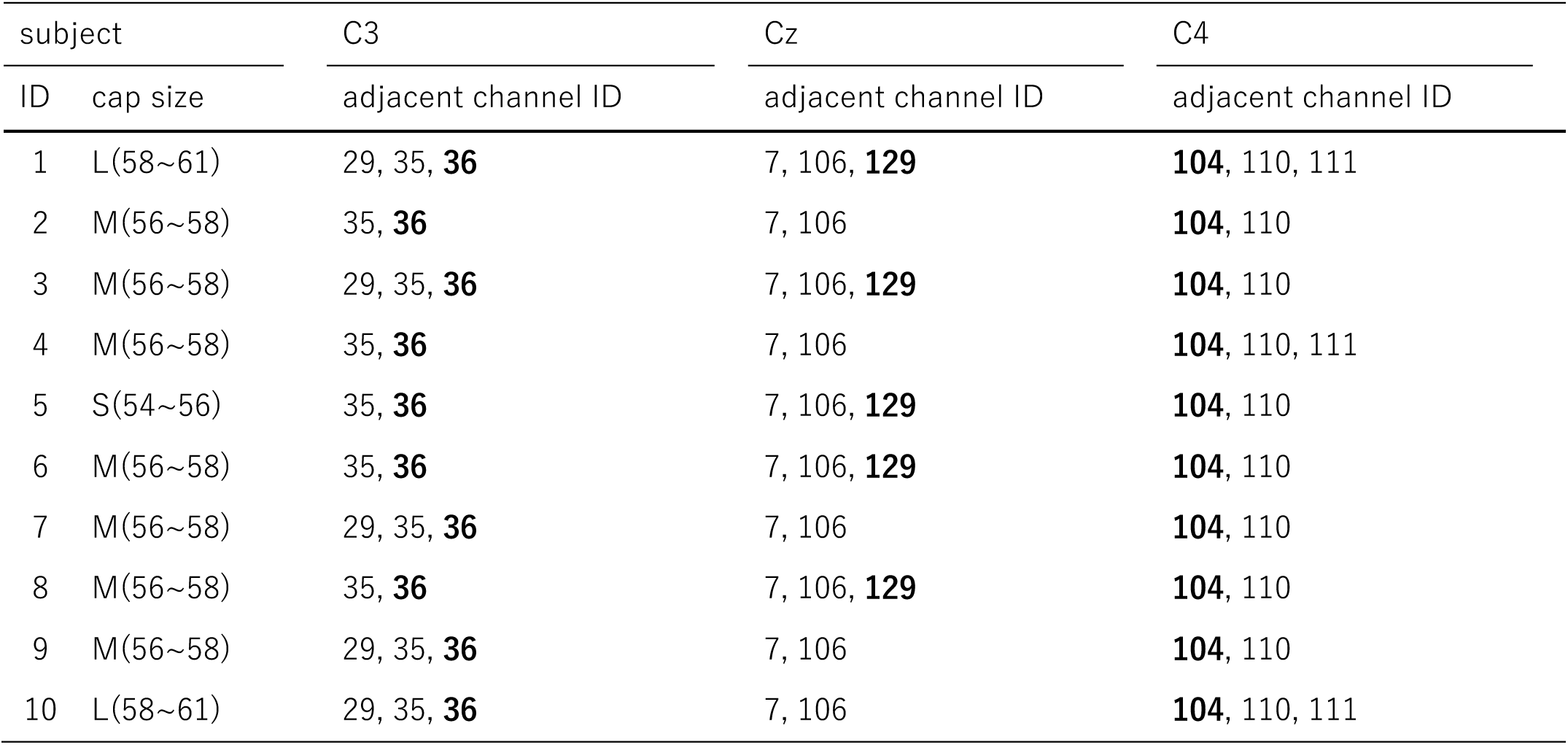
Electrode positions obtained with the proposed method. The cap size column describes the information on the HD-EEG net cap (Hydrocel Geodesic Sensor Net). C3, C4, and Cz electrodes in the 10-20 system are assigned for ID=36, 104,129 respectively.

| subject |  | C3 | Cz | C4 |
| --- | --- | --- | --- | --- |
| ID | cap size | adjacent channel ID | adjacent channel ID | adjacent channel ID |
| 1 | L(58~61) | 29, 35, **36** | 7, 106, **129** | **104**, 110, 111 |
| 2 | M(56~58) | 35, **36** | 7, 106 | **104**, 110 |
| 3 | M(56~58) | 29, 35, **36** | 7, 106, **129** | **104**, 110 |
| 4 | M(56~58) | 35, **36** | 7, 106 | **104**, 110, 111 |
| 5 | S(54~56) | 35, **36** | 7, 106, **129** | **104**, 110 |
| 6 | M(56~58) | 35, **36** | 7, 106, **129** | **104**, 110 |
| 7 | M(56~58) | 29, 35, **36** | 7, 106 | **104**, 110 |
| 8 | M(56~58) | 35, **36** | 7, 106, **129** | **104**, 110 |
| 9 | M(56~58) | 29, 35, **36** | 7, 106 | **104**, 110 |
| 10 | L(58~61) | 29, 35, **36** | 7, 106 | **104**, 110, 111 |

